# Surprising effects of stimulus repetition on neuronal firing rates and gamma-band synchronization in awake macaque V1

**DOI:** 10.1101/2025.07.09.663517

**Authors:** Martina Pandinelli, Mohsen Parto-Dezfouli, Eleni Psarou, Iris Grothe, Pascal Fries

## Abstract

Stimulus repetition is abundant, because the environment is redundant and/or because it is redundantly sampled. This offers an opportunity to optimize the processing of repeated stimuli. Indeed, stimulus repetition leads to classically described neuronal response decreases, and to more recently described neuronal gamma synchronization increases (sometimes preceded by decreases for a few trials). Here, we used a full-screen colored background (FSCB) and a flashed black bar, while recording multi-unit activity (MUA) and local field potentials (LFP) from area V1 of an awake macaque monkey. We found that the FSCB repetition induced neuronal response increases (sometimes preceded by decreases for a few trials) and gamma synchronization decreases (preceded by increases for a few trials). These effects are largely opposite to the dominant previous findings. Intriguingly, these surprising effects reversed when we isolated the responses to the flashed black bar. We discuss these findings, considering differences to previous studies with regards to the subject of the study, the stimuli and the task. We notice that in studies reporting classical results for gamma, sometimes in combination with firing rates, the stimuli were typically (partly) predictive of the reward. Here, we found non-classical results for the FSCB that was not reward predictive, and classical results for the black bar that was reward predictive. Whether this has revealed a general effect of reward predictive versus non-predictive stimuli will require further investigation with stimuli and task designs tailored specifically for this question.

**Significance:** Natural visual experience often entails repetitions of the same stimulus. This allows optimization of the brain’s processing of those stimuli. Indeed, repeated stimuli typically induce decreasing firing rates and increasing gamma-band synchronization in many visual areas. However, in this study in awake macaque primary visual cortex, we report surprising opposite effects: under some conditions, stimulus repetition can result in neuronal firing rate increases and gamma-band synchronization decreases. We argue that these effects might relate to the fact that the repeated stimulus was not predictive of a reward. In fact, when we isolated the responses to a second stimulus that was added later in each trial and that was predictive of reward, the effects reverted again to the typical pattern.

## Introduction

Repetition is a key feature of visual experience on several levels. On a behavioral level, the eyes of a subject tend to perform return saccades, as a strategy to optimize foraging (Psarou et al., 2025; Wilming et al., 2013), and thereby revisit the same objects. This brings the same stimuli repeatedly into the receptive fields (RFs) of a given neuron or neuronal group in retinotopically organized visual cortical areas, like primary visual cortex, V1. On a stimulus level, a natural visual scene can contain dynamic stimulus repetitions, when e.g. image elements move back and forth and thereby in and out of the mentioned RFs. On a combined stimulus and behavioral level, stimuli can contain spatial repetitions (Torralba & Oliva, 2003), like e.g. trees in a forest, or faces in a crowd, and the behavioral sampling of these repeated items can stimulate those RFs repeatedly with similar stimuli. All these types of stimulus repetition offer the brain the opportunity to optimize its computation (Olshausen & Field, 1996; Schwartz et al., 2007).

Repetition of visual stimuli in experimental contexts has generally led to decreases in neuronal firing rates (Desimone, 1996; Li et al., 1993) and in the fMRI BOLD (functional magnetic imaging blood oxygenation level dependent) response (Grill-Spector et al., 2006). This effect, classically described as repetition suppression, has no detrimental impact on behavioral performance (Grill-Spector et al., 2006). Maintained performance might be achieved by neuronal synchronization (Gotts, 2003; Gotts et al., 2012) which might increase the impact of the synchronized neurons by means of feedforward coincidence detection (Salinas & Sejnowski, 2001) or entrainment (Fries, 2015). Indeed, increased neuronal gamma-band synchronization within and between awake macaque areas V1 and V4 has been reported as a result of stimulus repetition (N. M. Brunet et al., 2014). This repetition-related gamma increase has been confirmed by studies that additionally demonstrated stimulus specificity of the repetition effect, both in monkeys (Peter et al., 2021; Psarou et al., 2025) and humans (Stauch et al., 2021).

In the present study, we report surprising findings obtained serendipitously while analyzing the effect of stimulus repetition in a dataset that had originally been collected to further investigate a different previously reported phenomenon: visually induced gamma-band activity rhythmically modulates the gain of the spike response to a stimulus change (Ni et al., 2016). In the present experiment, we recorded multi-unit activity (MUA) and local field potentials (LFPs) from awake macaque area V1, while inducing sustained gamma-band activity with a full-screen-colored background (FSCB) (Peter et al., 2019; Shirhatti & Ray, 2018). On top of the FSCB and at an unpredictable time, we then presented in the RFs of the recorded V1 neurons, a black bar that evoked neuronal spike responses.

This paradigm differed in several aspects from previous studies that had described repetition-related changes in gamma-band activity. These previous studies had used sustained stimulation with gratings or natural stimuli (N. M. Brunet et al., 2014; Peter et al., 2021; Psarou et al., 2025; Stauch et al., 2021), whereas here we used the FSCB and a flashed bar. Furthermore, in those previous studies, the stimuli that induced the investigated neuronal activity underwent changes or offsets that temporally predicted the reward. In the present experiment, reward was tightly predicted by the black bar, whereas the FSCB did not add to reward prediction.

This paradigm produced repetition-related changes in neuronal activity that were surprising, as they differed drastically from the results of those previous studies. The previous studies had confirmed numerous reports that stimulus repetition leads to MUA decreases (N. Brunet et al., 2014; Peter et al., 2021; Psarou et al., 2025). Furthermore, they revealed that stimulus repetition can lead to initial gamma decreases for a few (typically 4-10) repetitions (Peter et al., 2021; Stauch et al., 2021), always followed by gamma increases for the remaining repetitions (N. M. Brunet et al., 2014; Peter et al., 2021; Psarou et al., 2025; Stauch et al., 2021). By contrast, the present study revealed that the repetition of the FSCB induced MUA increases, preceded by MUA decreases for some conditions and/or task period. The repetition of the FSCB consistently induced gamma increases for some initial trials followed by gamma decreases for the remaining trials. Even more surprisingly, the repetitions of the black bar induced MUA decreases and initial gamma decreases followed by gamma increases, largely following results from previous studies.

## Results

Stimuli and task are illustrated in Fig. 1A. After an inter-trial interval with a randomly colored noise mask, each trial started with the presentation of a gray FSCB and a central fixation point (we refer to gray as a color, simply for ease of use). As soon as the monkey attained fixation, the FSCB color remained gray in 10% of the trials (gray-condition), or turned red in 90% of the trials (red-condition). The sequence of gray- and red-condition trials was pseudorandom. After 1.4-1.7 s, while the FSCB remained unchanged, a black bar (width: 0.1 dva (degrees of visual angle), length: 0.3 dva) was flashed for 0.2 s into the previously mapped RF, while the monkey was required to keep fixation. In different trials, the bar was presented at different contrast levels, pseudorandomly chosen from 5%, 10%, 20%, 40%, 80%, or 100% in each trial of the red condition, and always at full contrast (100%) in the gray-condition (see Methods for definition of contrast). Thus, the trials of the red-condition contained approximately equal numbers of trials of the 6 distinct contrast conditions. If the monkey kept fixation, the bar was followed by a gray FSCB and reward.

**Fig. 1.**
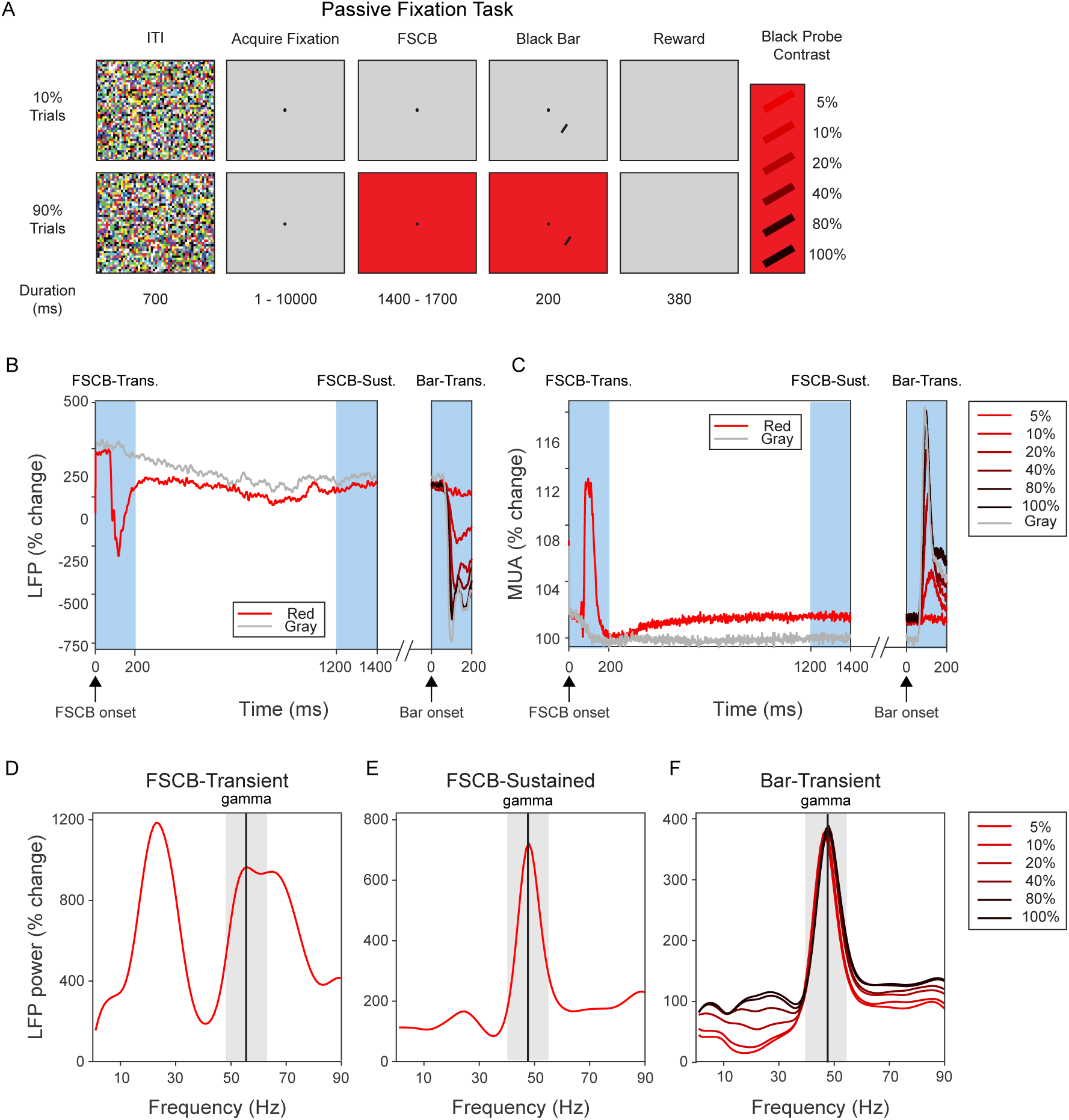
Stimuli, paradigm, task periods, and stimulus evoked and induced MUA and LFP responses. (*A*) Schematic illustration of stimuli and task periods for the gray condition (upper row) and red condition (lower row). (*B*) LFP averaged over trials and recording sites, separately for the gray and the red conditions, shown in the corresponding colors. (*C*) Same as *B*, but for MUA. (*D*) LFP power spectra during the FSCB-transient period for the red condition, averaged over recording sites and trials. Power is expressed as percentage change relative to the corresponding task period of the gray condition. (*E*) Same as *D*, but for FSCB-sustained period. (*F*) Same as D, but for bar-transient period. For the bar-transient period (B, C, F), the different bar-contrast conditions are color coded according to the legend to the right of panel *C*.

This paradigm offered an interesting opportunity to investigate repetition-related changes in neuronal activity, because it differed from previous studies as outlined in the introduction. We studied neuronal activity in response to the repetition of the FSCB and the repetition of the FSCB with the black bar superimposed. Moreover, we studied the neuronal response to the repetition of the black bar in isolation, by subtracting the response to the FSCB.

To analyze neuronal activity, we defined three periods indicated by the blue-shaded rectangles in Fig. 1B, C. The “FSCB-transient” period started with the fixation onset, which also triggered the red FSCB onset in the red condition; the FSCB-transient period was 200 ms long. The “FSCB-sustained” period corresponded to the last 200 ms end-aligned to the onset of the black bar. Finally, the “bar-transient” period started with the onset of the black bar and was 200 ms long. As expected, the LFP (Fig. 1B) and MUA (Fig. 1C) averaged over all trials and recording sites showed phasic responses during the FSCB-transient period in the red condition, much smaller phasic responses during this period in the gray condition (as there was no color change, just fixation onset), and contrast-dependent phasic responses in the bar-transient period.

For each of the conditions and periods, we first computed the raw LFP power spectra averaged over recording sites and all trials (Fig. S1). The raw LFP power spectrum is not very sensitive for showing rhythmical neuronal synchronization (Vinck et al., 2013). Nevertheless, these raw power spectra revealed gamma peaks or at least gamma bumps for all combinations of conditions and task periods. Note that clear gamma peaks were visible also for the FSCB-transient and FSCB-sustained periods of the gray condition. This was unexpected, and potentially due to a color-aftereffect, as few gray-condition trials were interspersed between many red-condition trials. We nevertheless opted to use the gray condition as a baseline condition for the normalization of the red condition. This is mainly useful for the investigation of power spectra, which are otherwise dominated by a 1/f component. Importantly, when we analyzed the raw LFP power spectra, all results remained qualitatively identical.

For the normalization, we expressed the power during the red condition as percent change relative to the gray condition (from the corresponding task period). For the bar-transient period, this was done separately per contrast condition. This was first done per recording site, and the resulting LFP power change spectra were then averaged over recording sites. They showed prominent beta and gamma peaks for the FSCB-transient period (Fig. 1D), and they were dominated by a large gamma peak for both the FSCB-sustained period (Fig. 1E) and the bar-transient period (Fig. 1F). For the bar-transient period, they differed across contrast conditions, mainly in the alpha-beta range (Fig. 1F). For further analysis of the gamma-band power, we defined the gamma band, separately per task period, as the peak frequency in the average power change spectrum plus/minus 10% of that frequency, resulting in 55±5.5 Hz for the FSCB-transient period, 48±4.8 Hz for the FSCB-sustained period, and 47±4.7 Hz for the bar-transient period (as indicated by the thin vertical lines and gray rectangles in Fig. 1 D-F).

### Repetition-related changes in MUA

We investigated repetition-related changes in MUA across trials. We first normalized all MUA values per recording site and per task period, by expressing MUA as percent of the MUA in the gray condition, averaged over the respective task period. These normalized MUA values were subsequently averaged over recording sites and plotted per trial (Fig. 2). We separately considered all combinations of color conditions (Fig. 2 rows) and task periods (Fig. 2 columns). Note that the bar-transient period during the red condition contained six different contrast conditions. Therefore, we also compiled a plot of repetition-related MUA changes, for which we averaged the data first per contrast condition and then over contrast conditions (Fig. 2D).

**Fig. 2.**
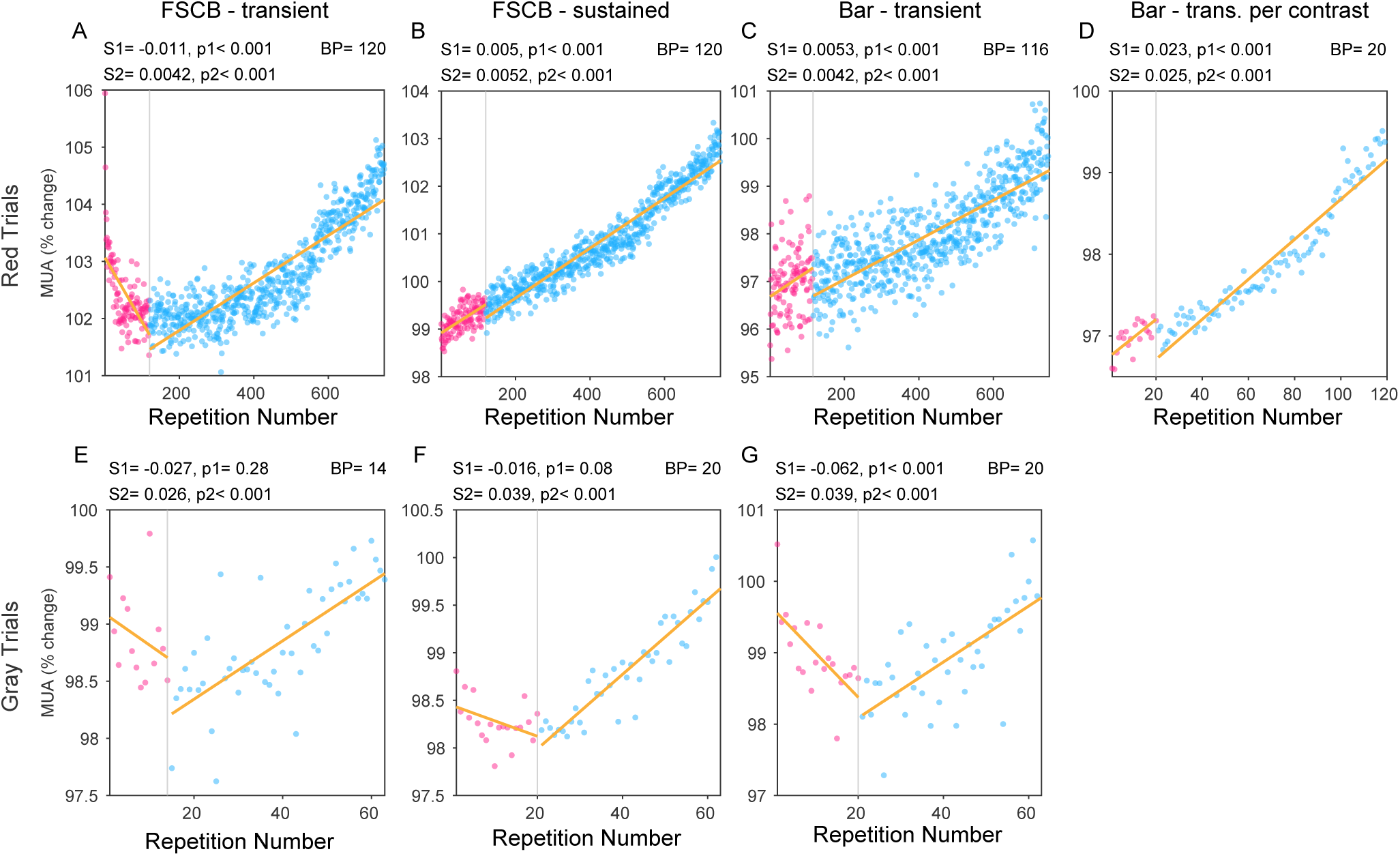
Trial-by-trial analysis of repetition-related changes in MUA responses. (*A*) MUA responses for the FSCB-transient period of the red condition, expressed as percent change to the corresponding gray condition, averaged over recording sites and sessions, separately per trial. Repetition-related changes were fitted with a segmented linear regression, whose results are shown as yellow regression lines, separated by a vertical line at the break point. The text above the plot provides the trial number of the break point (BP), and the slope and p-value for the first regression segment (S1, p1) and the second segment (S2, p2). The respective data points were colored magenta and blue. (*B*) Same as *A*, but for the FSCB-sustained period. (C) Same as *A*, but for the probe-transient period. (*D*) Same as *A*, but for the probe-transient period after averaging separately per bar-contrast condition. (*E*) Same as *A*, but for the FSCB-transient period of the gray condition. (*F*) Same as *A*, but for the FSCB-sustained period of the gray condition. (*G*) Same as *A*, but for the bar-transient period of the gray condition.

For several of the combinations of color condition and task period, MUA showed an initial repetition-related decrease followed by an increase. Similar breakpoints had been previously observed by us in several other datasets, both for MUA and gamma in awake macaque V1 (Peter et al., 2021; Psarou et al., 2025) and for gamma in human MEG (Stauch et al., 2021). In these datasets, the breakpoints had been determined by eye and occurred at trial 4-10 for gamma and 4-20 for MUA. In the current dataset, visual inspection of the MUA results suggested that breakpoints could occur around trial 20 when we considered the gray condition (Fig. 2E-G), and around trial 120 when we considered the red condition (Fig. 2A). We noticed that the red condition contained 6 different contrast conditions for the bar that was flashed into the RF, whereas the gray condition contained only 1 contrast condition. We further noticed that the breakpoint appeared around trial number 20 times the number of contrast conditions. We hypothesize that this might be related to the fact that the bar was temporally predictive of the reward (see Discussion for details). With this in mind, we opted for a segmented regression, where we fixed the number of segments to two, and the breakpoint to occur between trial 5-20 times the number of contrast conditions. This indeed provided excellent fits. For consistency, we applied this approach also to the combinations of color condition and task period, for which visual inspection did not reveal a breakpoint (Fig. 2B-D).

Most importantly, this analysis revealed that after the initial MUA decrease, which occurred for some conditions and task periods, all conditions and task periods showed repetition-related MUA increases until the end of the session. This was opposite to many previous reports, including our own, of repetition-related MUA decreases (N. M. Brunet et al., 2014; Peter et al., 2021; Psarou et al., 2025; Stauch et al., 2021).

We also investigated those repetition-related MUA changes as a function of time within the three task periods. For this, we focused on the trials after the breakpoint, and we pooled over red and gray conditions. After pooling, per task period, we determined the breakpoint (Fig. S2) and divided the remaining trials into three equal-sized trial bins, the early, middle and late trials (Fig. 3). Within each trial bin, we averaged the normalized MUA (blue, gray and red lines in Fig. 3A-C), and we statistically compared the late to the early trials (Fig. 3D-F). This showed that repetition-related MUA increases occurred throughout all three task periods.

**Fig. 3.**
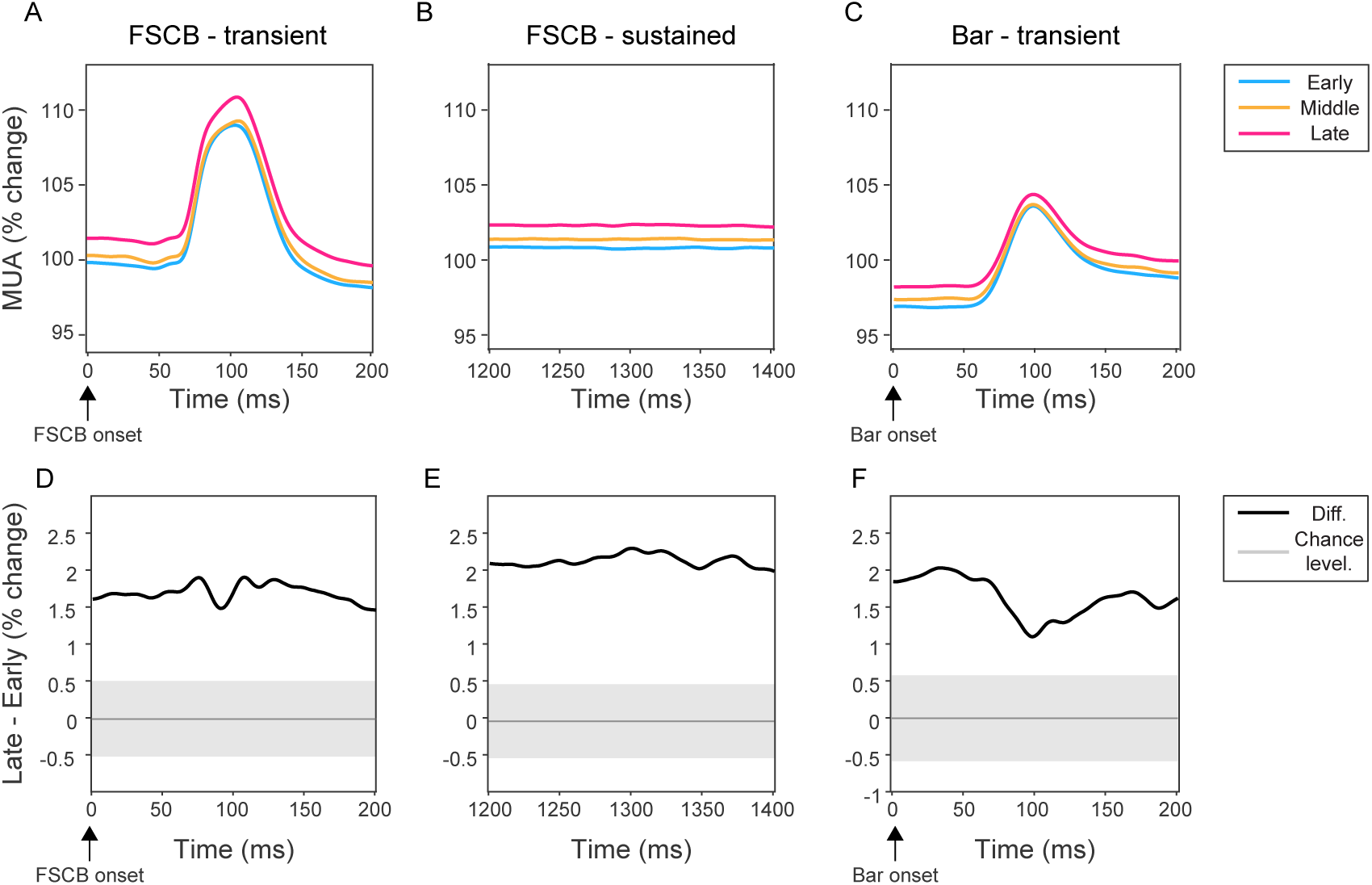
Time-course analysis of repetition-related changes in MUA responses. (*A*) MUA responses, expressed as percent change relative to the average FSCB-sustained period of the gray condition, for the FSCB-transient period of both the red and gray condition combined, separately for the early, middle and late trial group as indicated by the color legend right of panel *C*. The trial groups were defined after excluding the trials before the break point identified by the segmented linear regression. (*B*) Same as *A*, but for the FSCB-sustained period. (*C*) Same as *A*, but for the bar-transient period. (*D*) The black line shows the difference of the late minus the early trial group in *A*. The gray-shaded region indicates the range of differences expected by chance. All observed differences were significant. (*E*) Same as *D*, but corresponding to panel *B* for the FSCB-sustained period. (*F*) Same as *D*, but corresponding to panel *C* for the bar-transient period.

### Repetition-related changes in LFP gamma power

We next investigated repetition-related changes in LFP gamma-band power, using the same approaches as used for MUA. Gamma showed initial repetition-related increases followed by decreases, consistently for all combinations of color condition and task period (Fig. 4A-F) except for the bar-transient period in the gray condition (Fig. 4G). The segmented regression located the breakpoint to trial number 8-16 for the gray condition with one bar-contrast condition (Fig. 4E-G), at trial number 7 when we averaged over bar-contrast conditions in the red condition (Fig. 4D), and at trial number 30-50 for the red condition, which corresponds to trial number 5-8.3 after dividing by the 6 bar-contrast conditions (see last paragraph for the rationale of this). These breakpoint positions were consistent with the hypothesis that the breakpoint occurs at trial number 5-15 times the number of bar-contrast conditions.

**Fig. 4.**
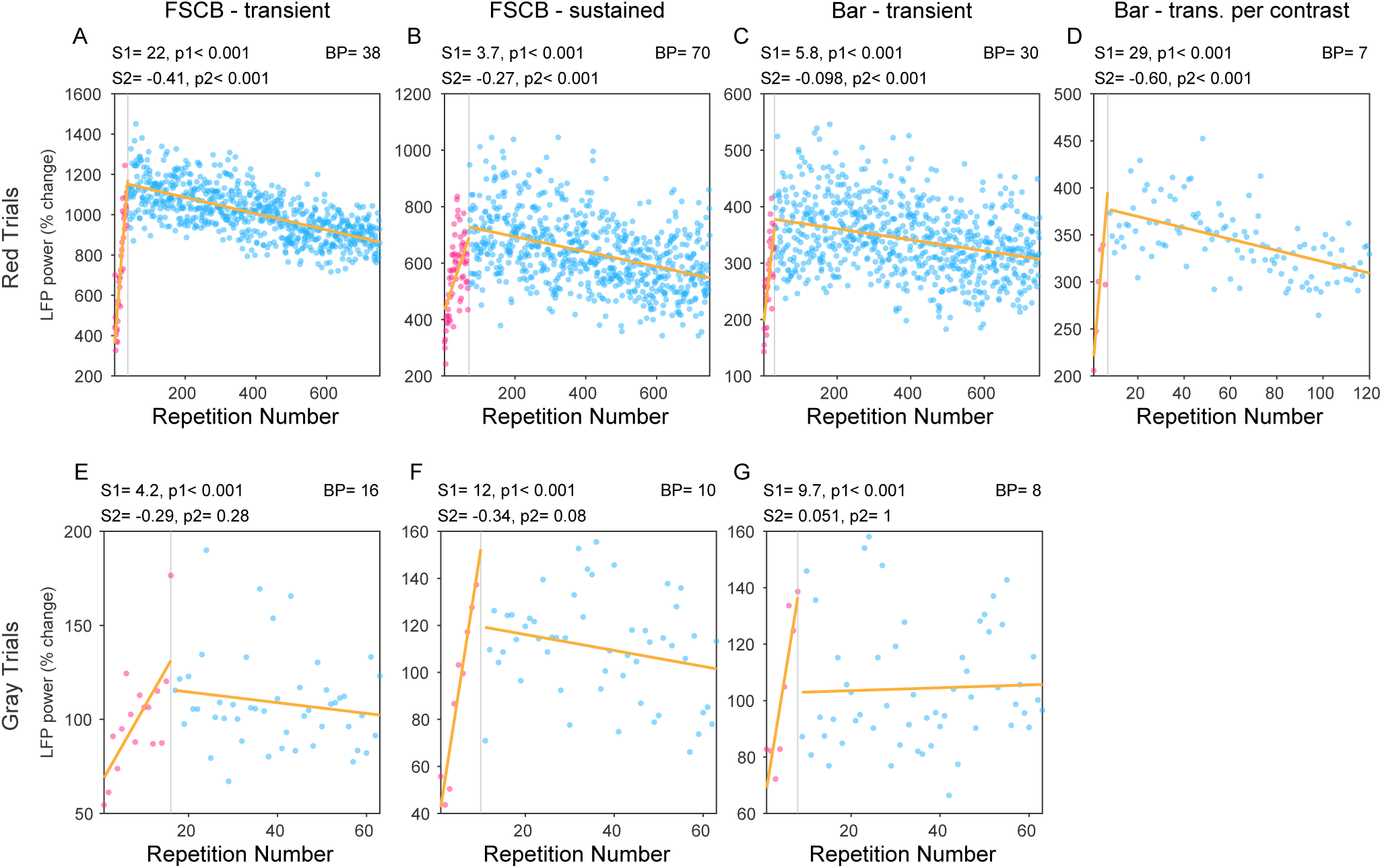
Trial-by-trial analysis of repetition-related changes in LFP gamma-band responses. Same as Fig. 2, but for LFP gamma-band power instead of MUA responses.

Most importantly, the observation of repetition-related gamma decreases preceded by initial increases is opposite to our previous studies using other stimuli and task paradigms. In those previous studies, we had consistently found repetition-related gamma increases, preceded in some cases by initial decreases for 4-10 trials.

We also investigated those repetition-related LFP-power changes as a function of frequency within the three task periods, following the same approach as for MUA (for trial-pooled gamma results and breakpoints see Fig. S3). This revealed, for the FSCB-transient period, repetition-related LFP power increases in the alpha band and decreases in both the beta and gamma bands (Fig. 5A, D). For the FSCB-sustained and the bar-transient period, repetition-related changes were dominated by gamma decreases (Fig. 5B, C, E, F). Note that both the FSCB-transient and the bar-transient period most likely contained event-related potentials (ERPs), both for the red and the gray conditions. As we used the gray condition for normalization, the ERP-related spectral perturbations might have influenced the normalized spectra. Therefore, we also considered an alternative normalization, in which the different task periods of the red conditions were always normalized by the FSCB-sustained period of the gray condition. This left the results essentially unchanged (Fig. S4).

**Fig. 5.**
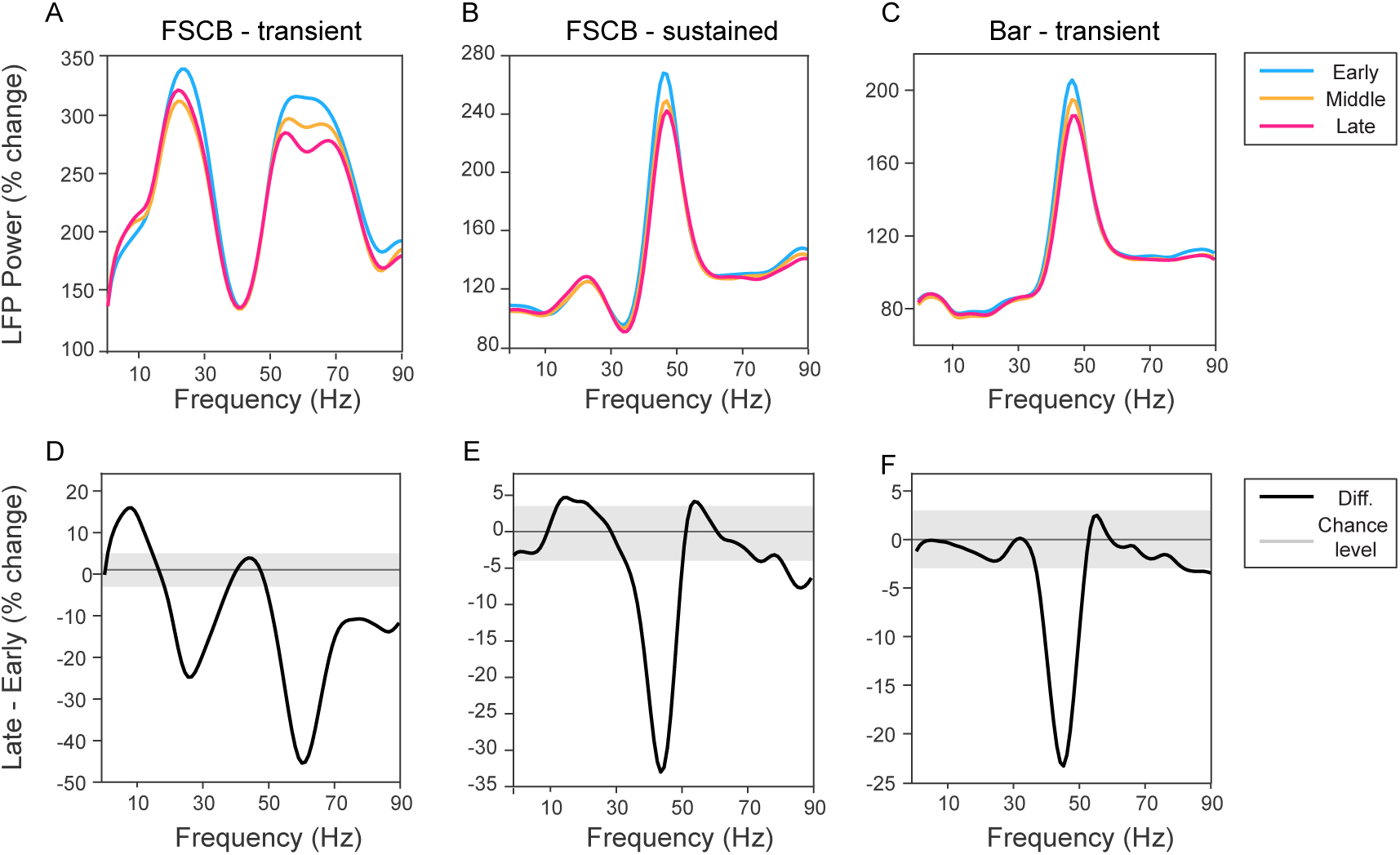
Spectral analysis of repetition-related changes in LFP power. Same as Fig. 3, but for LFP power changes during the different task periods, expressed as percent change relative to the corresponding task periods of the gray condition.

### Repetition-related changes in MUA-LFP gamma synchronization

We also investigated MUA-LFP synchronization, quantified by the MUA-LFP pairwise phase consistency metric (Vinck et al., 2010). As the PPC is poorly defined for single trials, we used the same trial-binning approach as in Figs. 3 and 5.

During the FSCB-transient (Fig. 6A, D) and the bar-transient periods (Fig. 6C, F), MUA-LFP PPC spectra were dominated by low-frequency synchronization, most likely due to the transient neuronal responses jointly evoked in MUA and LFP by the FSCB and bar onset, respectively. Those low-frequency components were the only that showed repetition-related changes, namely an increase during the FSCB-transient period and a decrease during the bar-transient period.

**Fig. 6.**
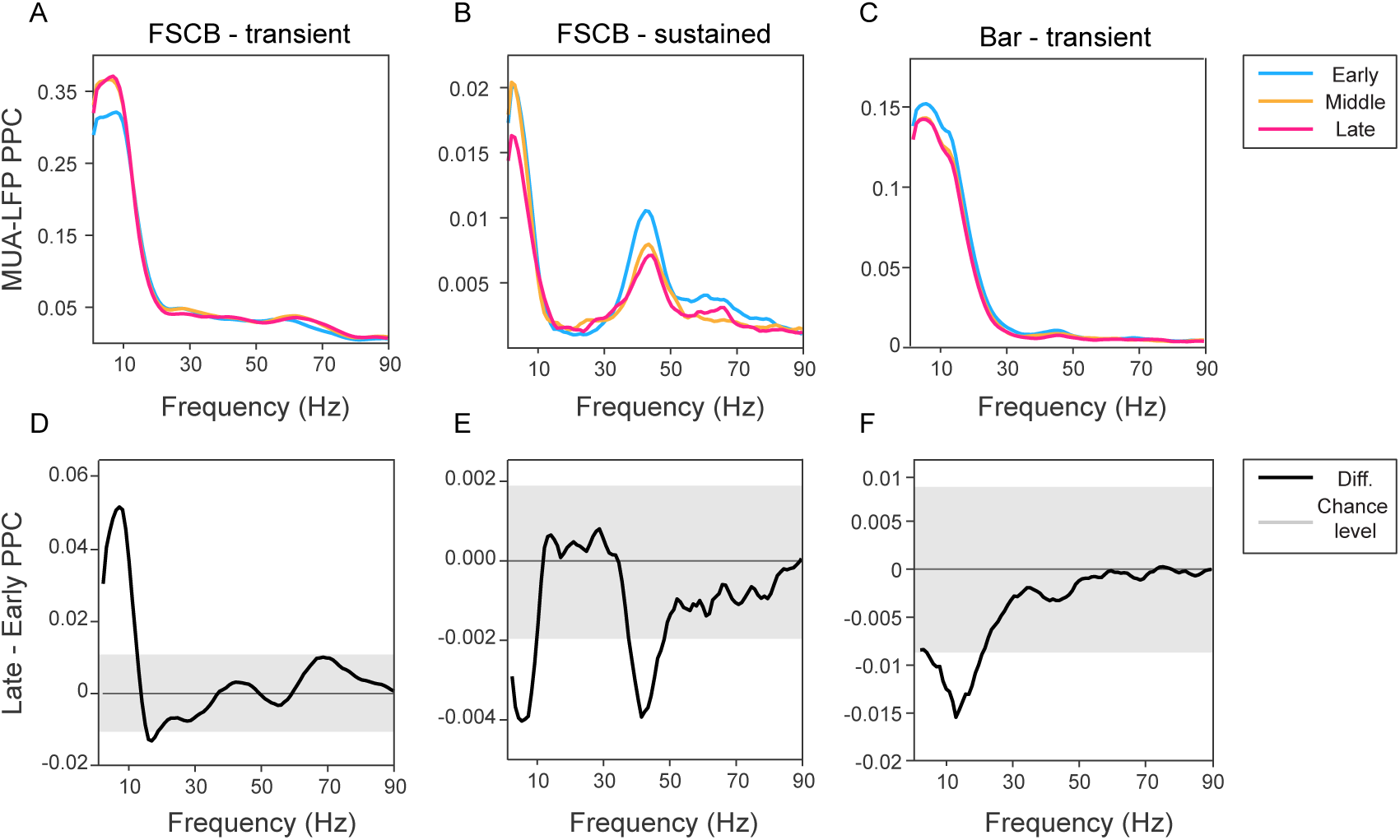
Spectral analysis of repetition-related changes in MUA-LFP PPC. Same as Fig. 3, but for MUA-LFP PPC spectra, which are intrinsically normalized.

During the FSCB-sustained period (Fig. 6B, E), the MUA-LFP PPC spectrum showed a low-frequency and a gamma component. Both showed repetition-related decreases. The MUA-LFP PPC gamma component and its repetition-related decrease corresponds closely to the gamma component in the LFP power spectra, suggesting that the LFP power spectra reflect local neuronal synchronization.

### Isolating responses to the bar and their repetition-related changes

As mentioned above, the repetition-related changes observed for the current stimuli and task were largely opposite to the changes observed previously (N. M. Brunet et al., 2014; Psarou et al., 2025; Stauch et al., 2021). Those previous studies had used gratings or photos of objects to induce neuronal activity, and those stimuli underwent changes or offsets that temporally predicted the reward. In the present study, the stimulus that temporally predicted reward was the bar. We have already analyzed the neuronal responses to the bar on top of the FSCB. Yet, this response to the compound FSCB + bar stimulus might be dominated by the response to the FSCB, and we therefore set out to isolate the response to the bar by subtracting the response to the FSCB alone. As a good estimate for this isolated FSCB response, we used the FSCB-sustained period. We subtracted the FSCB-sustained results from the bar-transient results, and refer to the difference as bar-only results (Fig. 7). We analyzed this separately for all red-condition trials pooled (Fig. 7A, D), for the red-condition trials averaged per bar-contrast condition (Fig. 7B, E), and for the gray trials (Fig. 7C, F). To our great surprise, those bar-only results showed again the same repetition-related changes that we had found in previous studies, namely for MUA a monotonic decrease (see Fig. 7A-C for significances), and for gamma an initial decrease followed by an increase for the rest of the trials (Fig. 7D-F, both highly significant for most conditions).

**Fig. 7.**
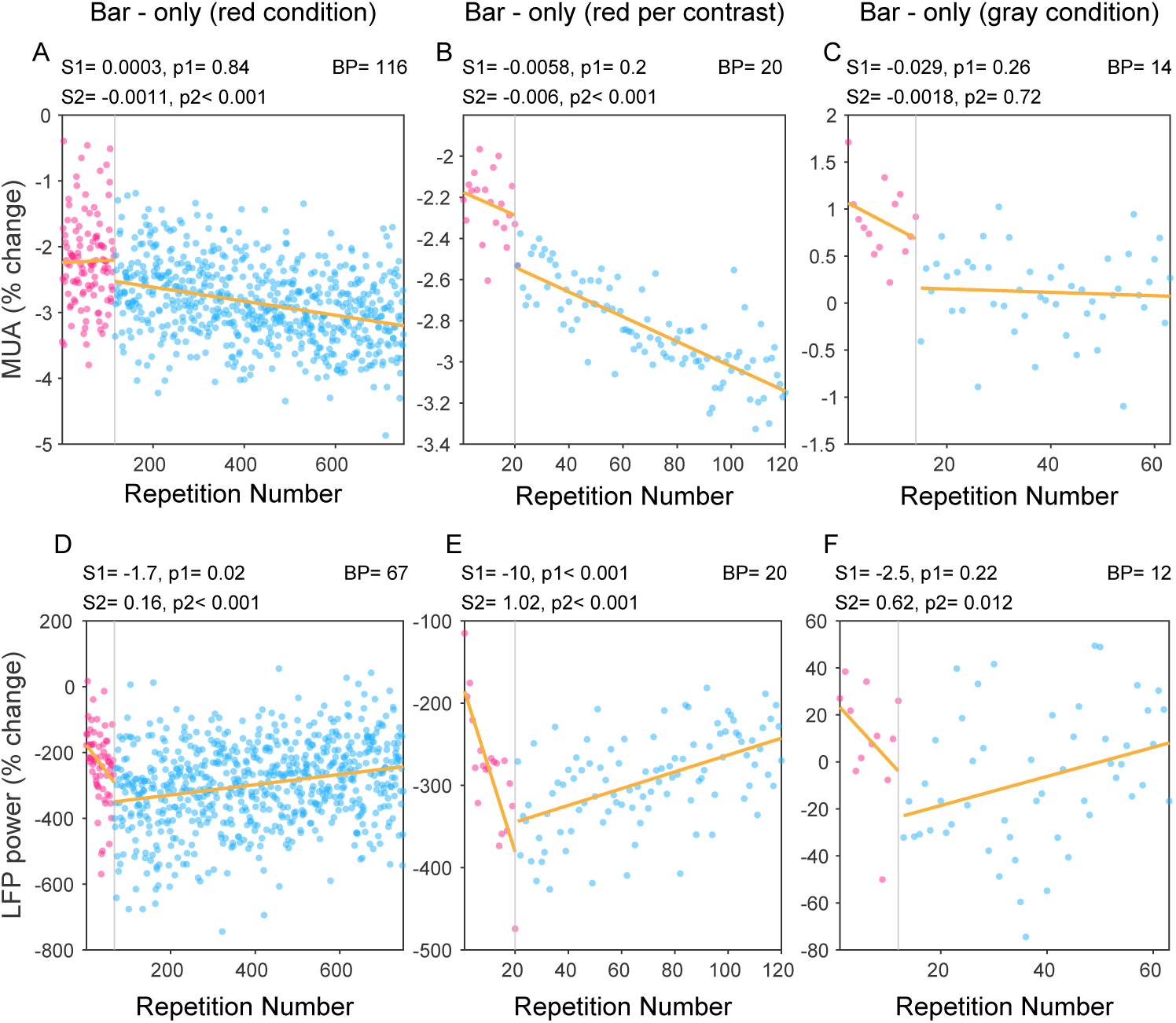
Isolation of MUA (upper row) and LFP gamma power (lower row) responses during the bar-transient period after subtracting the responses to the FSCB-sustained period. (*A*) Same as Fig. 2C, but after subtracting the data from the FSCB-sustained period. (*B*) Same as Fig. 2D, but after subtracting the data from the FSCB-sustained period. (*C*) Same as Fig. 2G, but after subtracting the data from the FSCB-sustained period. (*D*) Same as Fig. 4C, but after subtracting the data from the FSCB-sustained period. (*E*) Same as Fig. 4D, but after subtracting the data from the FSCB-sustained period. (*F*) Same as Fig. 4G, but after subtracting the data from the FSCB-sustained period.

### Correlations between repetition-related and stimulus-driven changes

Finally, we tested whether repetition-related changes in neuronal activity were correlated with stimulus-driven changes in neuronal activity. MUA in V1 can be enhanced by many stimuli, and the repetition of those stimuli typically causes MUA response reductions. By contrast, we found, for all conditions except the bar-only condition, repetition-related MUA response increases. Intriguingly, the MUA response in awake macaque V1 to large uniform red surfaces had previously been reported to be a reduction below baseline (Peter et al., 2019). If this stimulus-induced MUA reduction were itself reduced by stimulus repetition, just like stimulus repetition typically reduces stimulus-induced MUA enhancements, then this might explain our surprising observation of repetition-related MUA increases. If so, the repetition-related MUA increases should be negatively correlated with stimulus-induced MUA changes: the larger the stimulus-induced MUA decrease, the larger the repetition-related MUA increase. We quantified stimulus-induced MUA changes as the ratio of the red condition over the gray condition, and refer to this as the stimulus-related response ratio (SR). We quantified repetition-related MUA changes as the ratio of the last 30 trials over the first 30 trials, and refer to this as the repetition-related response ratio (RR). RR values were positively correlated to SR values during all task periods (Fig. 8A-C). This analysis also revealed that the stimuli in this dataset mostly did not suppress but enhance MUA, leading to positive SR values. Both of these observations do not support the possibility that the repetition-related MUA increases were due to repetition-related reductions in stimulus-induced response reductions.

**Fig. 8.**
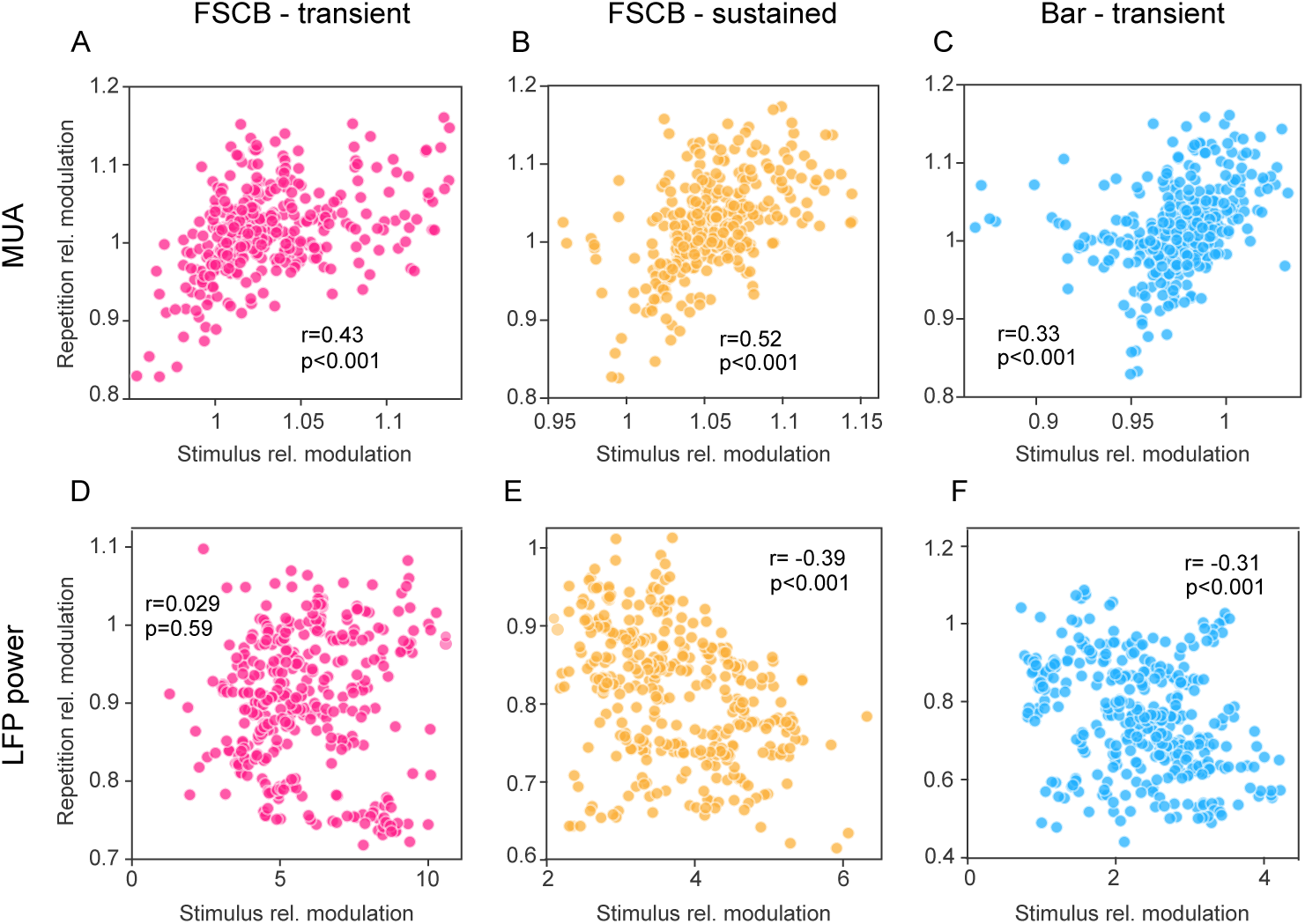
Analysis of correlation between repetition-related and stimulus related modulation. In all plots, each dot represents one recording site, the x-axis value represents the site’s stimulus-related modulation (stimulation-over-baseline ratio), and the y-axis value represents the site’s repetition-related modulation (late-over-early ratio). The upper row shows the MUA data, the lower row shows the LFP gamma power data. The three columns show the results separately for the three task periods, as indicated on the top. The insets in each plot provide the respective Pearson correlation coefficients and the corresponding p-values.

We repeated the same analysis for gamma-band power. For gamma, RR values were not correlated to SR values during the FSCB-transient period (Fig. 8D), and they were negatively correlated during the FSCB-sustained and bar-transient period (Fig. 8E, F).

## Discussion

### Summary

We report surprising serendipitous findings from investigating repetition-related changes in MUA and neuronal gamma-band activity in a new dataset. Such repetition-related changes have been reported in several previous publications, whose overall results can be summarized as follows: When a given stimulus is repeatedly presented, (1) MUA responses decrease monotonically with stimulus repetition, with a slope that is steep for the initial ∼20 trials and then flattens, (2) neuronal gamma-band responses increase with stimulus repetition, and this increase can be preceded by a decrease for the initial 4-10 trials. These repetition-related effects were largely reversed in the present dataset: (1) MUA responses decreased for the initial trials in some of the conditions or task periods, but subsequently, MUA responses increased for the remaining trials; (2) neuronal gamma-band responses consistently increased for the initial trials in both the red and the gray conditions and all task periods, and subsequently, neuronal gamma-band responses decreased for the remaining trials for all red-condition task periods and showed the same trend for the gray-condition task periods. However, when we isolated the bar-only response by analyzing the bar-transient minus the FSCB-sustained period, this largely reverted the effects back to those found previously.

### Limitations of the study

These surprising findings have been obtained from an experiment that investigated a different effect, namely gamma-rhythmic gain modulation (Ni et al., 2016), and this latter purpose determined stimuli and task. This experimental design leaves several possible explanations for the surprising findings, and it will require future experiments to arbitrate between them. Yet, the observations reported here and in our previous studies on repetition-related gamma changes do allow to argue against some and in favor of some other, intriguing, explanations, and we will spend most of this discussion on this exercise.

Note that also our original report of repetition-related gamma increases was based on a serendipitous finding in a dataset that had been obtained to study selective visual attention (N. M. Brunet et al., 2014).

The surprising findings are most likely due to experimental differences compared to previous studies. The main differences that appear plausible are: (1) Differences between animals, (2) differences between stimuli, (3) differences between tasks.

### An effect of the specific animal?

The animal used in the present study is different from the ones used in (N. M. Brunet et al., 2014) and (Peter et al., 2021), but the same as used in one previous study. This latter study with this same animal showed repetition-related changes in MUA and gamma-band responses that were highly similar to the ones reported in (N. M. Brunet et al., 2014) and (Peter et al., 2021). Thus, the specific animal used in the present study is likely not the reason for the surprising findings. Yet, this one previous study with this same animal also used similar stimuli and tasks as parts of (Peter et al., 2021) and (Stauch et al., 2021), leaving open the possibility that differences in the present study in either one of those aspects were relevant.

### An effect of the specific stimuli?

#### An effect of using uniform, colored stimuli?

The previous studies reporting repetition-related gamma increases had used gratings (N. M. Brunet et al., 2014; Peter et al., 2021; Stauch et al., 2021), and one of them had additionally used photos of natural objects (Peter et al., 2021). None of them had used uniform, colored surfaces. Thus, this specific choice of stimuli might appear like a plausible candidate to explain the surprising findings. However, we had six recording sessions from the same animal, in which we used uniform colored discs in a task that was highly similar to (Psarou et al., 2025). While this was a very limited dataset, the results were fully in line with our previously published reports (Fig. S5) and do not support an explanation of the surprising findings as a result of using these stimuli.

#### An effect of using full-screen stimuli?

The previous studies reporting repetition-related gamma increases had mostly used stimuli that did not cover the full screen. However, the original report demonstrated the effect first and very clearly for full-screen gratings (Fig. 1A-C of (N. M. Brunet et al., 2014)). Thus, the use of full-screen stimuli could clearly be eliminated as explanation for the surprising findings.

### An effect of the specific task?

This leaves the specific task as an intriguing candidate explanation. To explore this, we first review the main aspects of the tasks that have induced repetition-related gamma increases in previous studies.

In the original report of repetition-related gamma increases (N. M. Brunet et al., 2014), several different datasets were used. Most of the data were recorded while macaques performed selective attention tasks, during which two patches of drifting grating were presented and one was cued as behaviorally relevant. One of the patches was inside the RF of the recorded neurons, the other one outside, and attention was randomly cued per trial to the stimulus inside or outside. The cued stimulus had to be monitored for a randomly timed change in shape (for the ECoG data in that study) or color (for the V4 microelectrode data in that study), which the animal needed to report by releasing a bar. The behavioral response prompted the disappearance of both the attended and the non-attended stimulus, and their disappearance immediately preceded reward delivery. This same study also reported repetition-related gamma increases for another ECoG dataset. For this dataset, a monkey monitored the fixation point for a color change, while the neuronal activity was induced by a full-field patch of drifting grating. Similar to the attention tasks, the behavioral report of the fixation-point color change prompted the disappearance of the stimulus, which immediately preceded reward delivery.

In the subsequent study (Peter et al., 2021), macaques monitored stimuli for randomly timed changes, which were small localized contrast decrements randomly positioned on the stimuli. Those stimuli were either natural stimuli (color photos of isolated leaves, flowers, sweets, fruits, vegetables) or grating patches. Change detection was again reported manually, which prompted stimulus disappearance and subsequent reward delivery.

In the next study (Stauch et al., 2021), human subjects monitored a large grating patch for a suprathreshold (50%) contrast decrement of the entire grating combined with a near-threshold orientation change. The contrast change prompted the behavioral report of the orientation change. The behavioral report in turn prompted stimulus disappearance and subsequent presentation of a smiley, which provided positive feedback irrespective of behavioral response correctness.

In the most recent study (Psarou et al., 2025), macaques fixated while patches of grating were presented in the RFs of the recorded neurons. The duration of stimulus presentation was either 0.8-0.9 s for variable-stimulus blocks, or 1-1.3 s for fixed-stimulus blocks (see ref for details), and the stimulus offset was immediately followed by reward delivery.

In summary, in the previous datasets for which a repetition-related gamma increase has been reported (N. M. Brunet et al., 2014; Peter et al., 2021; Stauch et al., 2021), the stimulus that activated the recorded neurons was behaviorally relevant for obtaining the reward, and/or stimulus changes or stimulus offsets were predictive of the upcoming reward.

In the present study, as soon as the animal acquired fixation, the FSCB appeared and stayed on throughout the trial. After 1400 to 1700 ms, randomly timed, the bar was presented inside the RF of the recorded neurons for 200 ms. After the bar disappeared, the FSCB turned gray, which was a change for the red-condition trials and no change for the gray-condition trials, and this was followed by reward delivery after a constant interval of 100 ms. Thereby, the bar was predictive of the upcoming reward. Note that when we isolated the neuronal responses to the bar by subtracting the responses to the FSCB, we found similar repetition-related changes as in previous studies. In comparison, the FSCB was less predictive of the reward, as it was presented for a variable amount of time before reward delivery, and it also did not contain any temporal structure predictive of the upcoming bar and/or reward. For the FSCB, the main findings were opposite to the previous studies. Based on these considerations, we would like to speculate as follows.

We will first focus on the changes that we have observed for many repetitions. In the previous studies, that was the repetition-related firing-rate decrease and gamma increase; we will later discuss the changes observed in some datasets (including the present one) for the initial repetitions. Those many-repetition-related changes might be explained by a statistical dependence of the reward on the stimuli. When a stimulus repeatedly predicted a reward, we observed that the stimulus-induced neuronal responses showed firing-rate decreases and gamma increases. By contrast, and this is the main novel insight from the present study, when a stimulus was repeatedly presented but not reward predictive, or at least there was another stimulus that was much more tightly reward-predictive, then the stimulus-induced neuronal responses showed firing-rate increases and gamma decreases.

We next turn to the changes that we have observed for the initial repetitions in some of the previous studies. In those previous studies this was mainly a firing-rate decrease that was steeper than for the later repetitions, and a gamma decrease for the initial 4-10 repetitions preceding the later gamma increase. This gamma decrease was observed for non-overtrained natural stimuli in macaques and for non-overtrained gratings in human subjects (Peter et al., 2021; Stauch et al., 2021), and it was not observed for highly-overtrained gratings in macaques (N. M. Brunet et al., 2014; Peter et al., 2021; Psarou et al., 2025). Therefore, we have previously suggested that the initial presentations of non-overtrained stimuli induce an arousal reaction, which leads to increased gamma, and that this arousal and its enhancing effect on gamma decline over the course of few trials. In the present study, we find similar initial gamma decreases when we isolate the response to the bar, and we find opposite changes, i.e., initial gamma increases when we analyze the response to the FSCB.

In addition, we here make one intriguing observation, which holds both for the responses to the FSCB and the responses to the black bar: the number of those initial trials was similar to previous studies, i.e., in the range of 7 to 20 when we analyzed the gray condition, for which only one bar contrast was used, and when we analyzed the red condition after averaging over the different bar conditions. However, when we concatenated all trials of the red condition, the number of those initial trials was larger, in the range of 30 to 120. When this larger number was divided by 6, i.e., the number of bar-contrast conditions for the red condition, then we ended up with similar initial-trial numbers as in the other cases. This suggests that the changes observed for the initial trials “count” the number of repetitions of a particular black-bar. This might be related to the abovementioned, speculative, mechanism that results from the pairing of a particular stimulus with reward. Yet note, that this putative explanation departs from our previous interpretation of the initial-trial related changes as an arousal-related effect. The arousal in response to the FSCB is not expected to be modulated by the pairing of the preceding black bar with reward.

We emphasize that these latter paragraphs offer speculations intended to explain all observations as parsimoniously as possible, namely as the learning of a stimulus’ reward predictability. Yet, those observations have partly been made serendipitously – this applies to the original study (N. M. Brunet et al., 2014) and the present one – and therefore the stimuli and tasks were not optimized to investigate the observed phenomena, to minimize confounds, and to dissect different possibilities of explanation. At the same time, the observed effects are so striking and appear so fundamental, and the datasets from invasive recordings in awake non-human primates are so precious, that we think those serendipitous findings deserve to be reported. That said, the speculations offered as putative interpretations definitively require rigorous testing in other datasets and experiments that allow the dissection of different possibilities, and we anticipate that this will be a highly fruitful avenue for future research.

## Acknowledgments

This work was supported by DFG (FR2557/1-1, FR2557/2-1, FR2557/5-1, FR2557/7-1), EU (HEALTH-F2-2008-200728-BrainSynch, FP7-604102-HBP), NIH (1U54MH091657-WU-Minn-Consortium-HCP). We thank Dr. Alina Peter for the most useful scientific input, and Rasmus Roese for the invaluable day to day assistance and supervision in the laboratory work.

## Author contributions

Conceptualization: M.P., E.P., P.F.; investigation: M.P., E.P., I.G.; methodology: M.P., E.P., I.G., P.F.; software: M.P., M.P.D., E.P.; formal analysis: M.P., M.P.D., E.P., P.F.; writing – original draft; M.P., P.F. writing – review & editing, all authors; visualization: M.P., M.P.D., P.F.; supervision: P.F.; funding acquisition, P.F.

## Supplemental information

Document S1. Figures S1-S5.

---

**Fig. S1.**
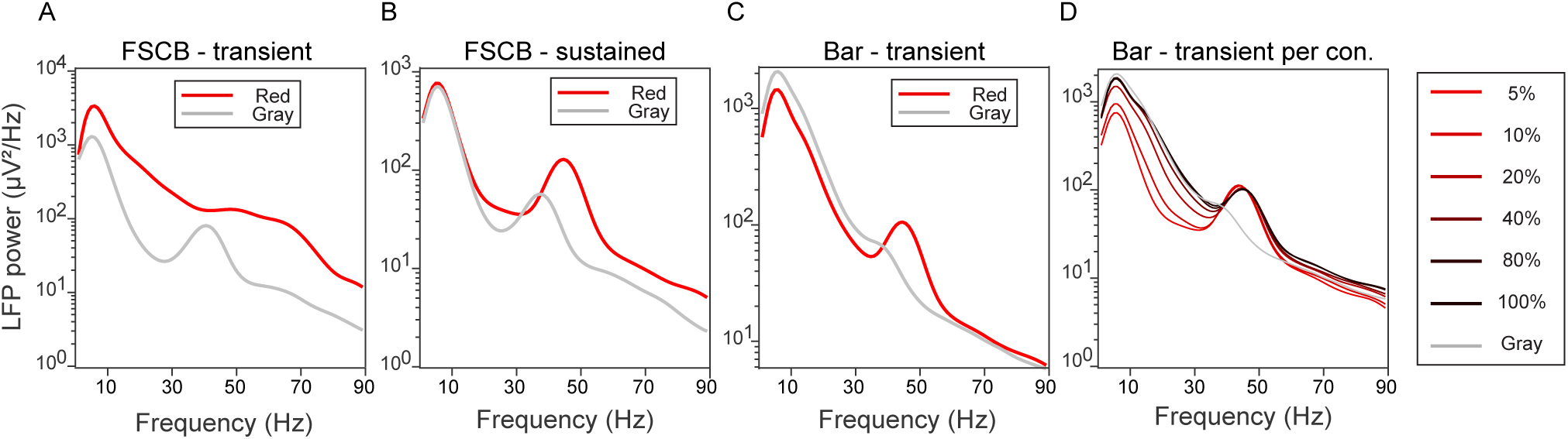
Raw LFP power spectra, without normalization. Each plot shows the raw LFP power spectrum, averaged over trials and recording sites, separately for the red and gray condition as indicated by the inset legends. The different plots show the data for the different task periods. For the bar-transient period, the data were either averaged over all contrast conditions (*C*), or separately per contrast condition (*D*), as indicated by the color legend to the right.

**Fig. S2.**
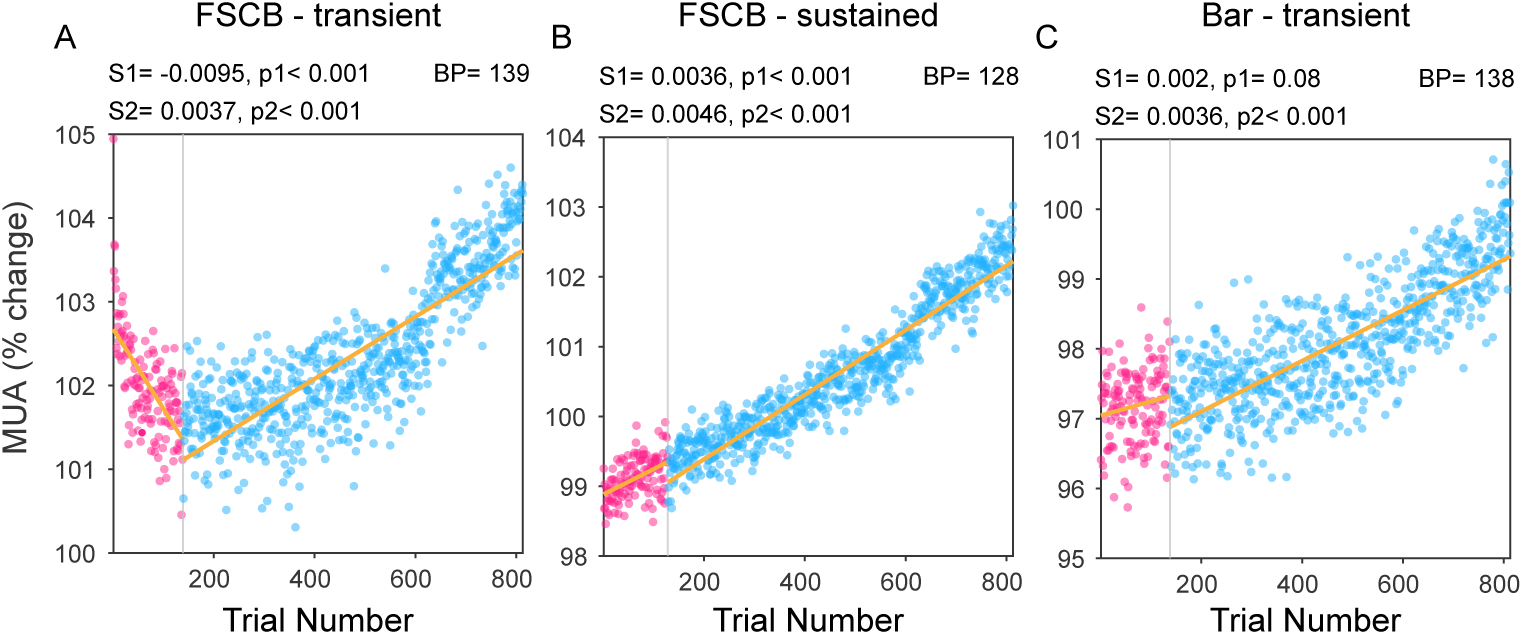
Same as Fig. 2, but for red- and gray-condition trials combined. As this combines trials of different conditions, those trials do not constitute repetitions of the same stimulus, and correspondingly, the x-axis is not labeled “Repetition number” but “Trial number”.

**Fig. S3.**
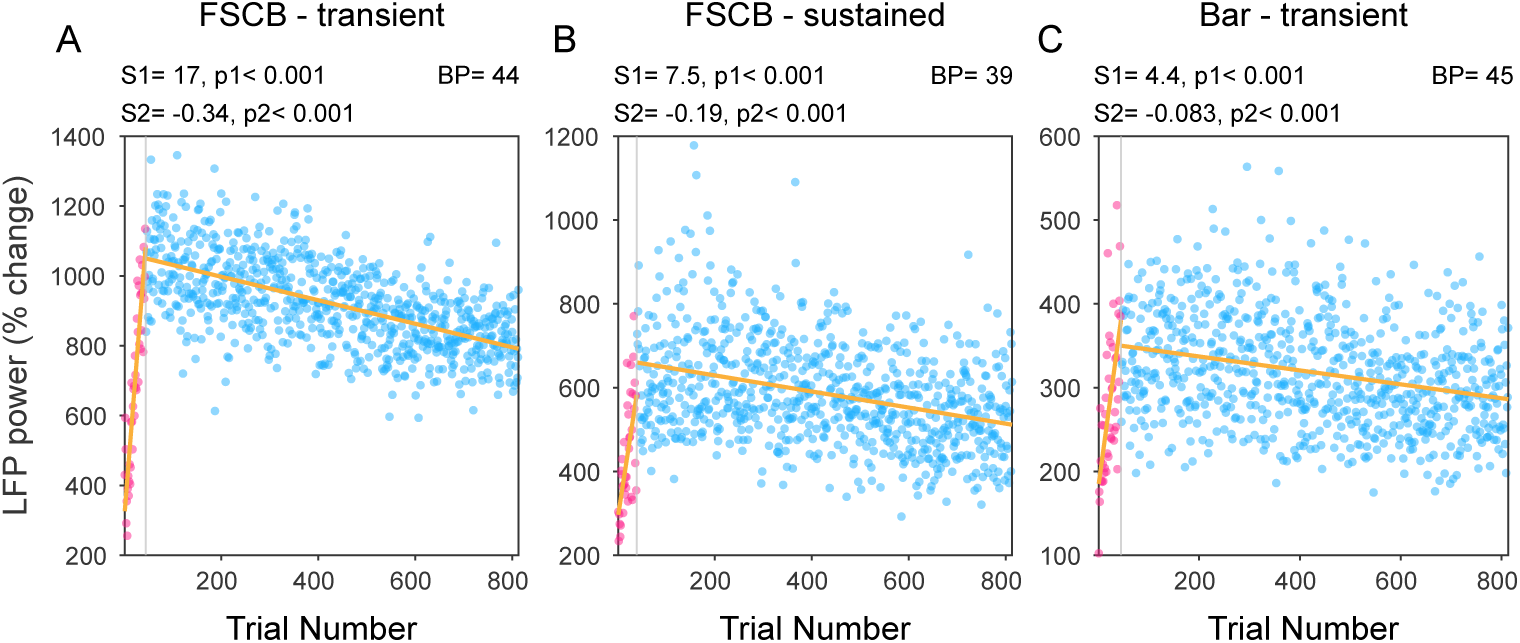
Same as Fig. 4, but for red- and gray-condition trials combined. As this combines trials of different conditions, those trials do not constitute repetitions of the same stimulus, and correspondingly, the x-axis is not labeled “Repetition number” but “Trial number”.

**Fig. S4.**
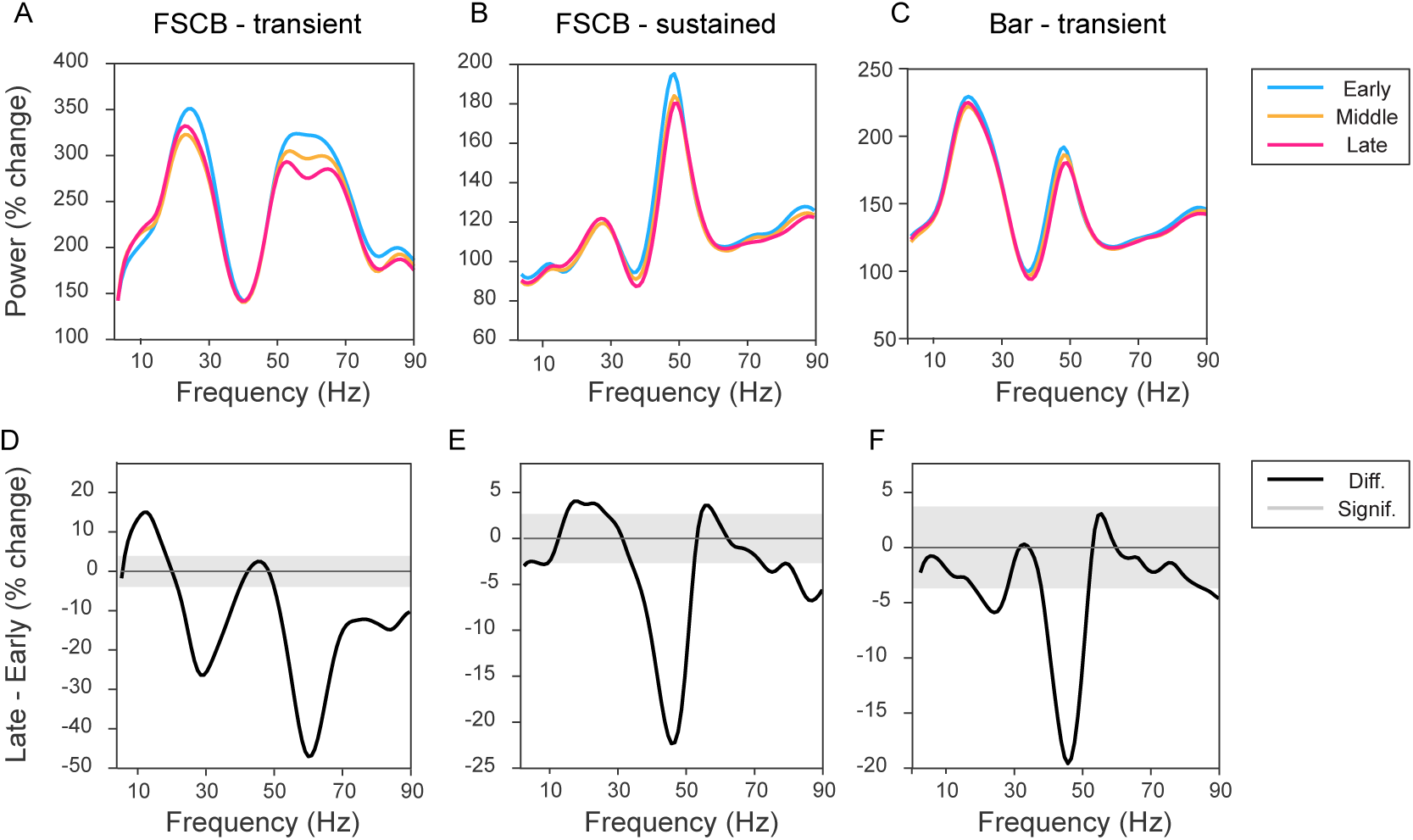
Same as Fig. 5, but always using the FSCB-sustained period for normalization.

**Fig. S6.**
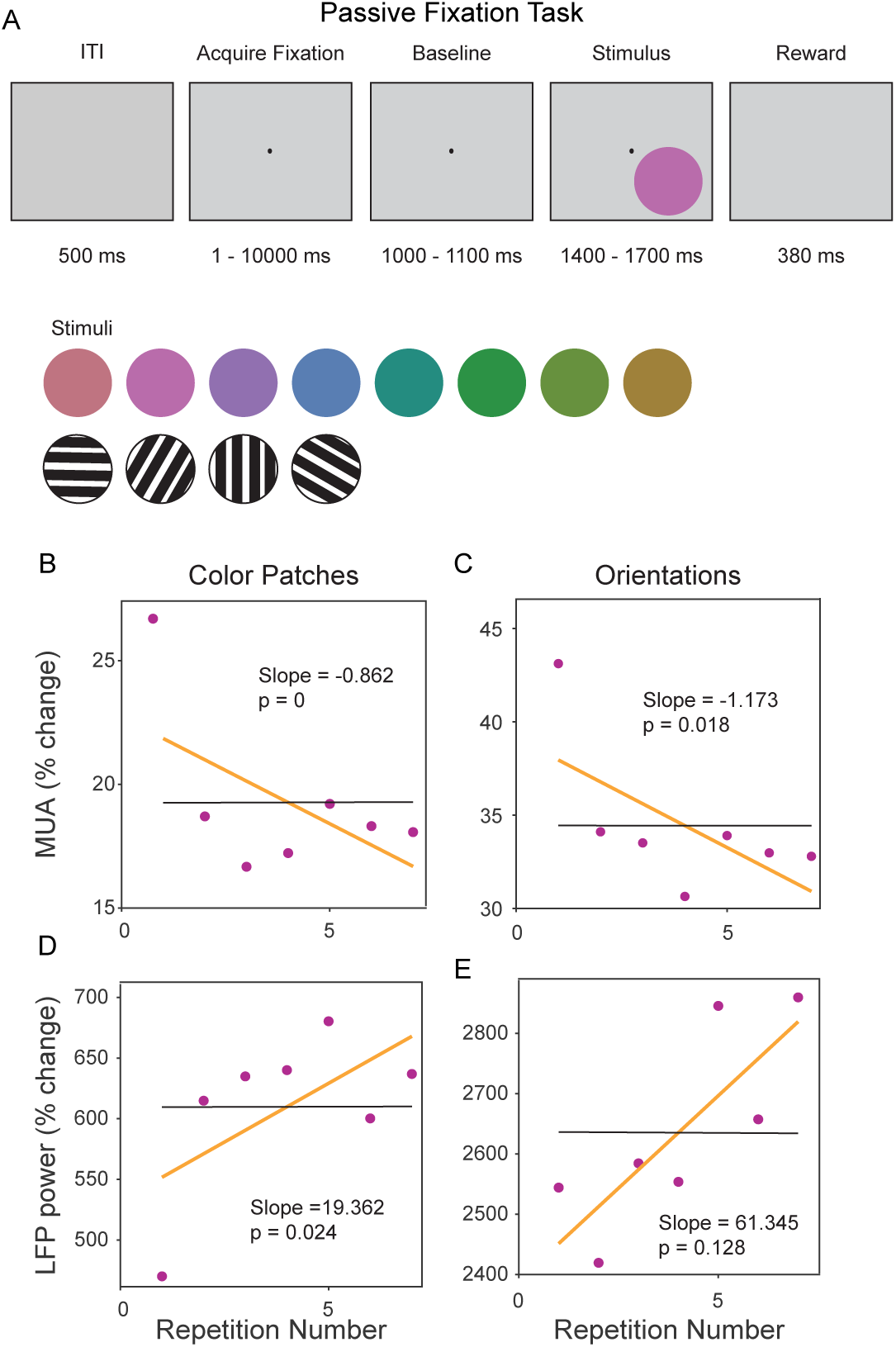
Classical results obtained in the same animal for different stimuli and task. (*A*) Illustration of task paradigm and stimuli. (*B*) Trial-by-trial analysis of repetition-related changes in MUA responses to the red-color disk stimuli. MUA responses for the stimulation period, averaged over recording sites and sessions, separately per trial. The yellow line shows the linear regression, whose slope and p-value are given as inset. (*C*) Same as *B*, but for grating patches. (*D*) Same as *B*, but for LFP gamma power. (*E*) Same as *C*, but for LFP gamma power.

## Materials and Methods

### Surgical procedures

All experiments on NHP were performed in compliance with the German and European ethical guidelines and regulations for the protection of animals. The experimental procedures were reviewed and approved by the German regional authority, the Regierungspräsidium Darmstadt.

One adult male macaque monkey (*Macaca Mulatta*), 19 years of age, weighing 13 kg, was used in this study. We used one rather than the more typical two or three animals, because we recently showed that the inference with the addition of a second or third animal is still limited to the sample (Fries & Maris, 2022; Laurens, 2022; Psarou et al., 2024). Thus, doubling or tripling the number of used animals would not change the quality of the inference, and ethical considerations mandate the use of one animal. All surgical procedures performed on the animal were carried out under general anesthesia, and analgesic and antibiotic treatments were administered in the postoperative time period.

The procedure for head-post implantation followed a two-step approach, described in detail in (Psarou et al., 2023). Subsequently, a modular custom-made titanium recording chamber for acute recordings was implanted. The chamber was fixed over the occipital area of the skull corresponding to visual area V1, on the left hemisphere. AAV virus injections into V1, used for other experiments, were performed at locations remote from the recording locations used here.

### Visual Stimulation

Visual stimuli were presented on an LG 32GK850G-B monitor (LG Electronics Inc., Seoul, South Korea), and gamma correction was applied in all tasks (gamma: 1.57). The monkey’s eyes were at a distance of 83.5 cm from the screen. The precise times of black-bar presentation were recorded by simultaneously presenting a small stimulus in the corner of the monitor, occluded from the monkey’s view, and recording that small stimulus with a custom-made photo-diode signal.

### Receptive field (RF) estimation

The RF position and orientation preference were determined after averaging neuronal responses over all electrode contacts on the laminar probe that were deemed to be located inside the cortex, as is explained in the corresponding section below. The electrode contacts were densely spaced, such that RF positions and orientation preferences were expected to be similar across simultaneously recorded contacts.

RF position was determined at the beginning of each recording session by using the back-projection method described in (Fiorani et al., 2014). Per trial, one bar moving in one of eight possible directions across the entire monitor was presented while the monkey kept fixation on a central fixation spot. The moving bar was 0.1 deg wide and had a speed of 8 deg/s. Ten correct trials for each direction were collected, for a total of 80 correct trials.

The MUA spike density functions were accumulated over time to reconstruct a spatial response map that indicated the position of the receptive field, after correcting for the response latency which maximized the back-projection response. The MUA’s response-based receptive field position was inferred by smoothing MUA responses with a Gaussian function for each direction of movement, and taking the 10th and 90th percentile and fitting an ellipse to the resulting data points.

### Orientation tuning task

The orientation preference of the located RF was inferred with a dedicated orientation tuning task. Per trial, a 0.1 degrees of visual angle (dva) wide and 0.3 dva long bar of one of six possible orientations, was presented at the location of the RF, while the monkey was required to keep fixation on the central fixation spot. A trial was considered correctly performed if fixation was maintained. Ten correct trials for each orientation were collected for a total of 60 trials. The preferred orientation was inferred by directly comparing the MUA (or LFP in case of low MUA signal) response amplitude across the 6 orientations and choosing the orientation with the highest related MUA response. The screen background during the orientation-tuning task was gray (RGB 128, 128, 128).

### Behavioral task

During training and recordings, the monkey sat in a primate chair located inside a fully shielded recording booth, with dim illumination. All recordings were performed under head fixation. The behavioral paradigm was created using ARCADE (Dowdall et al., 2018), a custom software built in house using MATLAB.

The monkey performed a fixation task. The animal acquired fixation on a central black fixation dot, 0.3 dva diameter, positioned in the center of the gray screen, and maintained fixation in a tolerance radius of 0.5 dva from the fixation spot. A trial was considered correctly performed when the monkey kept fixation for the whole duration of a trial (1650 to 1950 ms), and was rewarded with the delivery of grape juice.

Each trial consisted of the presentation of a sequence of visual stimuli on the monitor. Each trial began by showing a gray background screen (RGB coordinates [128, 128, 128]) together with the central fixation dot. The monkey was allowed a time period of maximally 10 seconds to acquire fixation, yet they typically attained fixation much quicker.

As soon as the animal started fixating, a full-screen colored background (FSCB) was shown. The FSCB was red colored (RGB coordinates [255,0,0]) in 90% of the trials (red-condition), while in 10% of the trials, the screen maintained the same gray color as the initial background (gray-condition). The gray trials were taken as a baseline reference for the red trials. The FSCB time period had a constant duration of 1450 ms, plus a time jitter ranging from 0 to 250 ms to precede the following section of the trial.

After the FSCB time period, the FSCB, red or gray, was kept in place, and an additional visual stimulus in the shape of a black bar was presented inside the previously determined RF for 200 ms. The black bar had a length of 0.3 dva, a thickness of 0.1 dva and was oriented according to the preference of the recorded neurons. In the red condition, the black bar stimulus was presented at 6 contrast levels (see below), while in the gray condition, the black bar stimulus was always shown at full contrast. After bar offset, a waiting period of 100 ms elapsed after which the animal was rewarded. Fixation breaks led to immediate termination of the trial, and no reward was given.

During each inter-trial interval (ITI, 700 ms), a colorful noise mask was presented on the screen. The noise mask was composed of squares of height and width ranging from 0.1 to 0.5 dva in steps of 0.01 dva, and each square had a color randomly selected from the possible RGB color coordinates. This mask was intended to counteract color adaptation effects.

### Color contrast and normalized color space

In the red condition, the black bar stimulus was presented at six contrast levels: 5%, 10%, 20%, 40%, 80%, 100%. The contrast was calculated based on the Weber contrast formula:

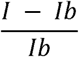

where *I* indicates the luminance of the stimulus, and *I_b_*indicates the luminance of the background. In order to define contrast levels appropriately, color coordinates for the red background and for the black bar foreground stimulus were chosen from the CIELAB standardized color space (Gravesen, 2015), then translated to RGB coordinates.

### Color patches and oriented gratings task

Results shown in Fig. S5 were obtained by analyzing data collected with a dedicated task. Every trial of the task started with an inter-trial interval (ITI) of 500 ms, presenting a grey screen (RGB [128,128,128]). An acquire-fixation time window started right after the ITI. A central fixation dot of 0.5 dva, dark gray in color (RGB [204,204,204]), was presented on the screen. In this time window, the monkey was allowed a maximum of 10000 ms to acquire fixation. After fixation was acquired, the monkey was required to maintain fixation for the duration of a baseline time window of 1000 – 1300 ms, during which the screen kept the initial grey color. At the end of the baseline window, the visual stimulus was presented during a 1300 – 1800 ms long time window. The stimulus consisted in a circular patch of uniform color or of grating. The circular-colored patch had a 7 dva diameter, covering RFs, and could assume one of the following colors: RGB [149,112,198], [188,107,180], [191,115,129], [160,130,60], [103,143,6], [44,147,59], [37,140,128], [90,126,179]. The grating stimulus was also circular, had a 3.5 dva diameter, a spatial period of 0.5 dva, and was approximately centered on the RF. The grating could assume one of four possible orientations: 0, 45, 90 or 135 degrees. If the monkey maintained fixation for the whole trial duration, the trial was considered correct, and the monkey was rewarded with delivery of grape juice.

### Eye position

Eye position was recorded at 1000 Hz using an Eyelink 1000 system (Eyelink Inc., SR research Ottawa, Canada). The system was calibrated on the left eye at the beginning of every recording session, and the left-eye signal was recorded.

### Recordings

In each recording session, one laminar probe was introduced through the intact dura into the cortex. Laminar probes featured 16 channels in one column, spaced by 50 micron (linear V-probes from Plexon Inc., Dallas, Texas, USA). The probe was guided by a guide tube that rested on the dura. For the recordings, the guide tube served as ground and the shank of the probe as reference. Recordings were performed in a fully shielded booth acting as a Faraday cage that prevented disturbance of the electrophysiological signal by line noise. Correspondingly, line-noise removal was not necessary and not performed.

Electrophysiological data were acquired using Tucker Davis Technologies (TDT, Alachua, Florida, USA) systems. Raw data were recorded and digitized by a TDT PZ2 pre-amplifier at 24.4140625 kHz.

### Data Analysis

All analyses were carried out in MATLAB and using the FieldTrip toolbox (Oostenveld et al., 2011).

#### Preprocessing

Raw data were downsampled to 1000 Hz. For the LFP, this entailed upsampling to 3,125,000 Hz, downsampling to 25,000 Hz, filtering with a stop-band at 500 Hz (IIR filter) and downsampling to 1000 Hz. For the MUA, this entailed upsampling to 3,125,000 Hz, band-pass filtering at 300-12000 Hz, rectification, stop-band filtering at 500 Hz (IIR filter) and downsampling to 1000 Hz.

#### Recording site selection

During each recording session, we performed one penetration with one laminar probe. The probe was positioned under online visual inspection of the neuronal signals and their general responsiveness to visual stimulation, such that typically all recording sites provided good MUA and LFP responses. Yet, some recording sites did not provide clear RFs, and those recording sites were identified by visual inspection and not used for further analyses. The remaining recordings sites will be referred to as “selected recording sites”, or “selected sites”.

#### Spectral analysis

For the spectral analysis, the LFP in the 200 ms long FSCB-transient, FSCB-sustained or bar-transient periods was multiplied with a Hann taper, zero-padded to 1s, and Fourier transformed using the FieldTrip function ft_freqanalysis.

#### Baseline normalization

All analyses except Fig. 1B, C and Fig. S4 use the gray condition as baseline, separately per task period. For the MUA, separately per recording site, each task period provided one value for the gray condition, averaged over time points and trials. This gray-condition value was used to express the corresponding red-condition MUA as percent change relative to the gray condition. The same was done for the LFP, except that the LFP was first transformed to a spectrum, and the averaging and normalization was done per frequency, providing a frequency-wise LFP power change spectrum during the red condition relative to the gray condition.

For Fig. S4, the same procedure was followed, except that the normalization was not by the corresponding task period of the gray condition, but always by the FSCB-sustained period of the gray condition.

For Fig. 1B, C the signals, both the raw LFP and the MUA, were expressed as percent change relative to the average respective values during the gray-condition FSCB-sustained period. This was done per recording site, with subsequent averaging over sites.

#### Estimation of gamma peak frequency and gamma band

The LFP power change spectra as compiled for the main analyses were used to determine the gamma peak frequency. The gamma peak frequency was estimated after averaging over all recording sites from all sessions, yet separately for the FSCB-transient, FSCB-sustained and bar-transient. The frequency associated with the maximum power change value in the 30-100 Hz range was defined as the gamma peak frequency, and a band of +/− 10% around the peak was defined as the gamma band.

#### Analysis of repetition-related changes across trials

To visualize and quantify repetition-related changes across trials, LFP gamma power and MUA were averaged over recording sites and sessions, separately per trial. For this, only the trials present in all sessions were used. In other words, the session with the smallest number of trials for the respective stimulation condition determined the number of trials that was used from all other sessions. When all trials of both the red and the gray condition were considered together, this resulted in 813 trials (Fig. S2 and S3). When all trials of the red condition were considered, this resulted in 750 trials (Fig. 2A, B, C, Fig. 4A, B, C, Fig. 7A, D). When all trials of the red condition were considered, yet averaged separately per bar-contrast condition, this resulted in 120 trials (Fig. 2D, Fig. 4D, Fig.7B, E). When all trials of the gray condition were considered, this resulted in 63 trials (Fig. 2E, F, G, Fig. 4E, F, G, Fig. 7C, F).

Visual inspection of the resulting trial-by-trial MUA and gamma values suggested that they could generally be described as a linear increase, or a linear decrease, or a combination of those two with a relatively well-defined break point in between. Furthermore, as discussed in the results and discussion sections, visual inspection suggested that the break point occurred at approximately trial number five to 20, times the number of bar-contrast conditions that was relevant for the respective analysis. Finally, we intended to treat all data as uniformly as possible. Therefore, we fitted the data with two linear regression lines, separated at a break point that was constrained to lie at trial five to 20, times the number of bar-contrast conditions that was relevant for the respective analysis. The number of bar contrast conditions that was relevant for the respective analysis was seven for the all-trials condition (Fig. S2 and S3), six for the red condition (Fig. 2A, B, C; Fig. 4A, B, C; Fig. 7A, D), one for the gray condition (Fig. 2E, F, G; Fig. 4E, F, G; Fig. 7C, F), and also one for the red condition after averaging over bar-contrast conditions (Fig. 2D; Fig. 4D; Fig. 7B, E). The decision about the precise trial constituting the break point was made objectively by using the approach described in (Bogartz, 1968). This approach is one version of a segmented linear regression. The algorithm iteratively considers each trial as a potential break point, and subsequently selects that trial as break point that minimizes the sum of the squared errors of the two regression lines.

#### PPC analysis

Local neuronal synchronization was assessed by calculating the Pairwise Phase Consistency (PPC) (Vinck et al., 2010) between MUA and the LFP (Fig. 6), as implemented in the FieldTrip toolbox (Oostenveld et al., 2011). For each session, respectively for each laminar-probe penetration, we calculated PPC between all combinations of MUA and LFP from all selected recording sites, except between MUA and LFP from the same recording site. This latter exclusion was made to minimize spectral bleeding of spikes into the LFP. We used the “unipolar” LFP, i.e., the LFP referenced against the laminar-probe shaft. Subsequently, the MUA-LFP PPC was averaged over site-pair combinations. PPC was based on Fourier transforms of the MUA and the LFP during the task period and trial groups as described in the results section.

#### Statistical analysis

Statistical testing was performed (1) for the segmented linear regressions quantifying the trial-by-trial MUA or gamma responses (Fig. 2, 4, 7, S2, S3), (2) for the simple linear regression quantifying the trial-by-trial MUA or gamma responses in the very few trials with color or grating patches (Fig. S5), (3) for the comparison of MUA change time courses, LFP change spectra, or MUA-LFP PPC spectra between late and early trial groups, including correction for multiple comparisons across time points or frequencies (Fig. 3, 5, 6, S4).

Regarding the statistical tests for case (1): The segmented linear regression was performed on the observed trial-by-trial data, giving two observed regression slopes. Subsequently, the trial numbers were randomized, and the segmented linear regression repeated, and this randomization was performed 1000 times. The slopes obtained across randomizations were collected in two randomization distributions, one for each segment. Per segment, we first determined whether the observed slope was positive or negative and then determined the number of randomization slopes with the same sign that exceeded the observed slope. This number was divided by the total number of randomizations and multiplied by two, and the resulting value was taken as p-value of a two-sided test.

Regarding the statistical tests for case (2): This followed the same approach as case (1), except that the regression was not segmented, but performed across all of the few available trials.

Regarding the statistical tests for case (3): We first obtained the observed difference time course or observed difference spectra of the late minus the early trial group. Subsequently, we randomly exchanged trials between the late and the early trial groups and recalculated the differences, and this was performed 1000 times. Importantly, for each randomization, only the minimal and the maximal difference – across all time points or frequencies - was retained and placed into the minimal randomization distribution and the maximal randomization distribution, respectively. The 2.5^th^ percentile of the minimal randomization distribution and the 97.5^th^ percentile of the maximal randomization distribution were taken as thresholds for statistical significance at a false-positive level of 5%, including correction for the multiple comparisons across time points or frequencies, respectively (Nichols & Holmes, 2002).

